# Inhibiting glycogen synthase kinase 3 suppresses TDP-43-mediated neurotoxicity in a caspase-dependant manner

**DOI:** 10.1101/2021.02.03.429569

**Authors:** Matthew A. White, Francesca Massenzio, Xingli Li, Michael P. Coleman, Sami J. Barmada, Jemeen Sreedharan

## Abstract

Amyotrophic lateral sclerosis-frontotemporal dementia (ALS-FTD) is a progressive and ultimately fatal disease spectrum characterised by 43-kDa TAR DNA-binding protein (TDP-43) pathology. Current disease modifying drugs have modest effects and novel therapies are sorely needed. We previously showed that deletion of glycogen synthase kinase-3 (GSK3) suppresses TDP-43-mediated motor neuron degeneration in *Drosophila.* Here, we investigated the potential of GSK3 inhibition to ameliorate TDP43-mediated toxicity in mammalian neurons. Expression of TDP-43 was found to both activate GSK3 and promote caspase mediated cleavage of TDP-43. Inhibition of GSK3 reduced the abundance of full-length and cleaved TDP-43 in rodent neurons expressing wild-type or disease-associated mutant TDP-43 and also ameliorated neurotoxicity. Our results suggest that TDP-43 turnover is promoted by GSK3 inhibition in a caspase-dependent manner, and that targeting GSK3 activity could have therapeutic value.

## Introduction

Amyotrophic lateral sclerosis (ALS) and frontotemporal dementia (FTD) are progressive and fatal neurodegenerative diseases that exist on a clinicopathological spectrum (ALS-FTD)[1]. Clinically, ALS is characterised by motor dysfunction, while FTD leads to a decline in cognition affecting executive functions, behaviour and language capabilities. The available disease-modifying drugs have only a minor impact on survival and disease progression, and novel therapeutic agents are therefore urgently required.

Almost all ALS and half of FTD cases are characterised by cytoplasmic ubiquitinated inclusions positive for TAR DNA-binding protein 43 kDa (TDP-43)[2–4]. Disease-linked mutations in *TARDBP* (the gene encoding TDP-43) indicate a fundamental role for TDP-43 in ALS-FTD pathogenesis[5–8]. TDP-43 inclusions are also seen in Alzheimer’s disease[9, 10], Parkinson’s disease[11, 12], Huntington’s disease[13] and limbic-predominant age-related TDP-43 encephalopathy (LATE)[14]. These observations implicate aberrant homeostasis of TDP-43 in a broad range of neurodegenerative diseases.

Caspases[15–18] and calpains[19, 20] can cleave TDP-43 to generate 25, 35 and 42 kDa C-terminal fragments. The accumulation of these phosphorylated and aggregated C-terminal fragments are a hallmark of ALS-FTD[3, 21, 22]. Cleavage products of TDP-43 are degraded by the proteasome and through autophagy[23–25]. However, whether aggregated and cleaved TDP-43 actually mediate disease or are non-toxic bi-products of physiological TDP-43 processing[26, 27] is unclear.

Glycogen Synthase Kinase-3 (GSK3) is a highly conserved and ubiquitously expressed serine/threonine protein kinase with wide ranging biological functions including glycogen metabolism, cell proliferation and apoptosis[28]. Mammalian GSK3 is encoded by two gene paralogues, *GSK3A* and *GSK3B,* which give rise to two protein isoforms GSK3α and GSK3β. Several lines of study link GSK3 biology to ALS-FTD pathogenesis. Firstly, expression of GSK3 is significantly increased in thoracic spinal cord tissue of patients with apparently sporadic ALS[29] and increased expression of GSK3 isoform β can be seen in frontal, hippocampal, cerebellar, cervical and lumbar tissue of patients with ALS or ALS with cognitive impairment[30]. Secondly, TDP-43 expression activates GSK3[31] whose activity modulates ER-mitochondrial associations regulated by vesicle-associated membrane protein-associated protein-B[32]. Thirdly, GSK3 is a modulator of TDP-43 cytosolic accumulation during cellular stress and its inhibition reduces the cytosolic accumulation of C-terminal TDP-43 fragments[33]. Finally, in an unbiased *in vivo* screen we previously showed that deletion of *shaggy,* the *Drosophila* orthologue of GSK3, significantly suppresses TDP-43-induced motor axon and neuromuscular junction degeneration[34]. Collectively, these data suggest that increased GSK3 activity plays a key role in neurodegeneration associated with TDP-43. Here, we confirm that GSK3 inhibition mitigates TDP-43-linked neurodegeneration in mammalian neurons and explore the biochemical and cellular mechanisms underlying this protective effect.

## Results

### TDP-43 activates GSK3 and undergoes caspase-mediated cleavage

To explore the mechanistic links between TDP-43 and GSK3 we began by expressing wild-type (TDP-43^WT^) and ALS-linked mutant (TDP-43^Q331K^) TDP-43 isoforms in human SH-SY5Y neuroblastoma cells. To control for nonspecific effects of transgene overexpression, we also transfected cells with an N-terminally truncated form of TDP-43 (TDP-43^N-Del^) (Fig 1a). TDP-43^N-Del^ has several attractive features as a negative control isoform as it lacks the region essential for dimerization and self-oligomerisation, which are critical steps necessary for many of the physiological functions of TDP-43 including nucleic acid binding [35]. N-terminal multimerization is also linked with the subcellular distribution of TDP-43 and its aggregation propensity[36–39]. All isoforms were C-terminally GFP-tagged (Fig 1a). Immunoblotting of cell lysates for GSK3 demonstrated that TDP-43^Q331K^ increased the activition of both GSK3α and GSK3β, as evidenced by reduced phosphorylation of serine 21 and serine 9 respectively. TDP-43^WT^ enhanced activation of GSK3α alone and TDP-43^N-Del^ had no significant effect on GSK3 activity (Fig. 1b,c). These observations are in keeping with previous studies demonstrating that TDP-43 activates GSK3[31].

**Fig.1.**
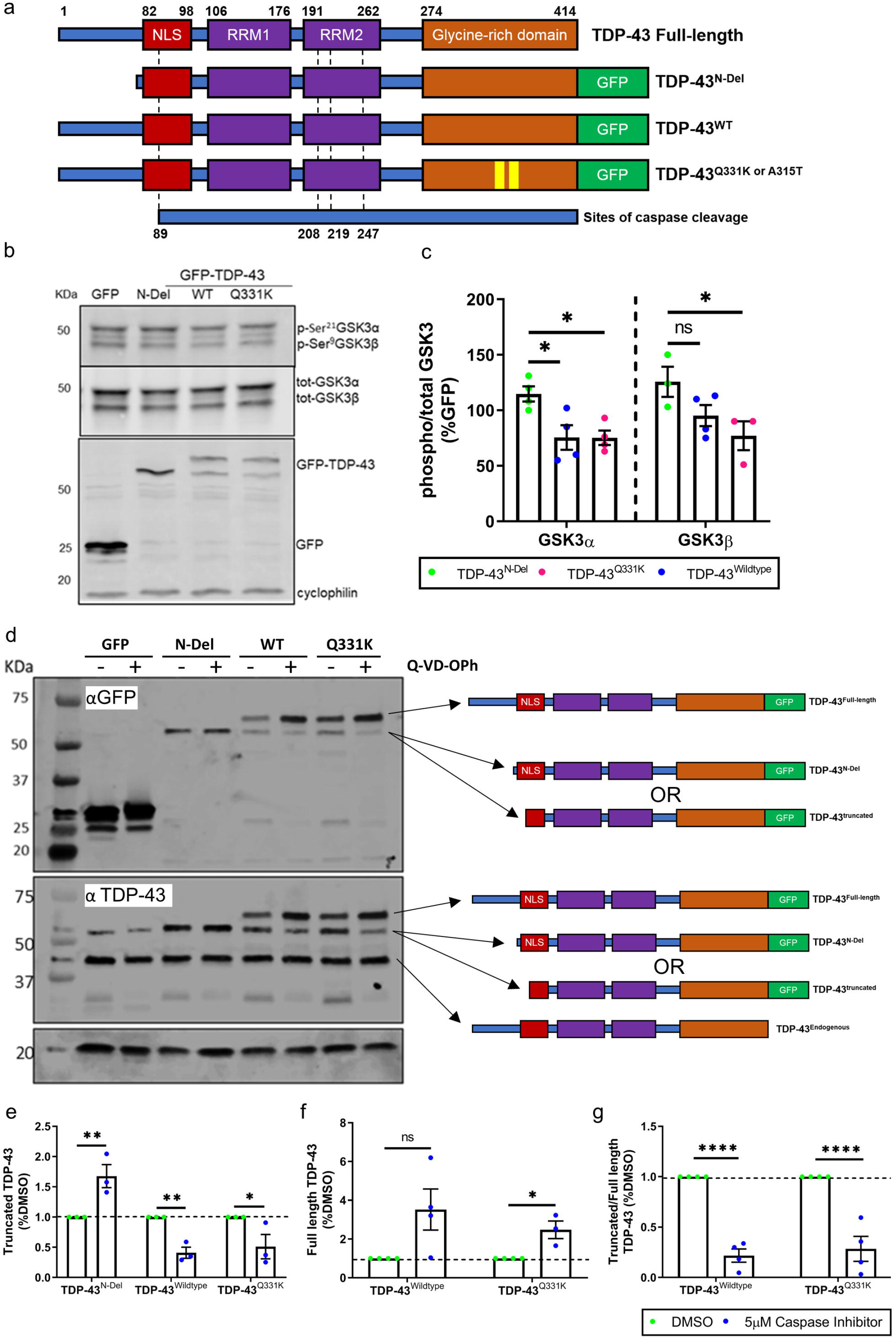
TDP-43 activates GSK-3 and undergoes caspase-mediated cleavage. **a.** Schematic of TDP-43 expression constructs used in this study. Dotted lines denote sites of caspase mediated cleavage of TDP-43. **b.** Representative immunoblot of SH-SY5Y cells transfected with expression constructs for GFP and GFP tagged: TDP-43^N-Del^, TDP-43^WT^ or TDP-43^Q331K^ for 24 hours and blotted for phospho-GSK-3α/β (Ser21/9) and total GSK-3α/β. **c.** Immunoblot band intensity quantification (*n* = 3-4 biological replicates per condition). The ratio between phospho-GSK3/total GSK-3 denotes GSK3 activation. ANOVA genotype *P* = 0.0012. Pairwise comparisons: GSK-3α, TDP-43^N-Del^ vs TDP-43^WT^: \**P*= 0.0186; TDP-43^N-Del^ vs TDP-43^Q331K^: \**P*= 0.0186; GSK-3β, TDP-43^N-Del^ vs TDP-43^WT^: ns *P* = 0.0515; TDP-43^N-Del^ vs TDP-43^Q331K^: \**P* = 0.0123. Two-way ANOVA followed by Holm-Sidak *post-hoc* test for pairwise comparisons. **d.** Representative immunoblot of SH-SY5Y cells transfected with expression constructs for GFP and GFP tagged: TDP-43^N-Del^, TDP-43^WT^ or TDP-43^Q331K^ for 24 hours with and without treatment with the pan caspase inhibitor Q-VD-OPh. **e-g.** Immunoblot band intensity quantifications showing the abundance of truncated, full length and the ratio truncated:full length TDP-43 in the absence of caspase activity (DMSO vs. Q-VD-OPh). **e.** Truncated TDP-43, pairwise comparisons: TDP-43^N-Del^: \*\**P* = 0.001679; TDP-43^WT^: \*\**P*= 0.004268; TDP-43^Q331K^: \**P* = 0.013091. **f.** Full length TDP-43, pairwise comparisons: TDP-43^WT^: ns *P* = 0.054872; TDP-43^Q331K^: \**P*= 0.011711. **g.** Ratio truncated:full length TDP-43. For **(e-g)** (n = 3-4 biological replicates per condition); ****P < 0.0001; multiple t test with Holm-Sidak correction for multiple comparisons. Error bars denote mean ± s.e.m.

We next immunoblotted the transfected cell lysates for both GFP and TDP-43. Interestingly, cells expressing TDP-43^WT^ or TDP-43^Q331K^ demonstrated two prominent bands corresponding to fulllength GFP-tagged TDP-43 and a smaller ~55kDa band, which was comparable in molecular weight to GFP-tagged TDP-43^N-Del^ (Fig. 1b, lower panel). As the TDP-43^N-Del^ construct was deliberately truncated near to a caspase cleavage site (Fig 1a), we hypothesised that the ~55kDa band seen after expression of full-length TDP-43 isoforms was a product of caspase cleavage. Indeed, application of a caspase inhibitor significantly reduced the abundance of the ~55kDa fragment and increased the abundance of full-length TDP-43^WT^ and TDP-43^Q331K^ (Fig. 1d-g). We concluded that overexpressed TDP-43 undergoes N-terminal caspase-mediated cleavage to generate C-terminal TDP-43 fragments.

### GSK3 inhibition preferentially reduces the abundance of truncated TDP-43

To explore the link between GSK3 activity and TDP-43 fragmentation, SH-SY5Y neuroblastoma cells expressing TDP-43^N-Del^, TDP-43^WT^ or TDP-43^Q331K^ were treated with the GSK3 inhibitors CHIR99021 and AZD1080 (Fig. 2a). Inhibition of GSK3 trends towards a reduction in the abundance of full length TDP-43^WT^ and TDP-43^Q331K^ (Fig. 2b). More strikingly, however, GSK3 inhibition significantly reduced the abundance of cleaved products of both TDP-43^WT^ and TDP-43^Q331K^ (Fig. 2c,d). Beyond cleaved products, GSK3 inhibition also reduced the abundance TDP-43^N-Del^ (Fig. 2c). These results suggest that a GSK3 mediated mechanism alters the abundance of caspase-cleaved TDP-43 C-terminal fragments.

**Fig.2.**
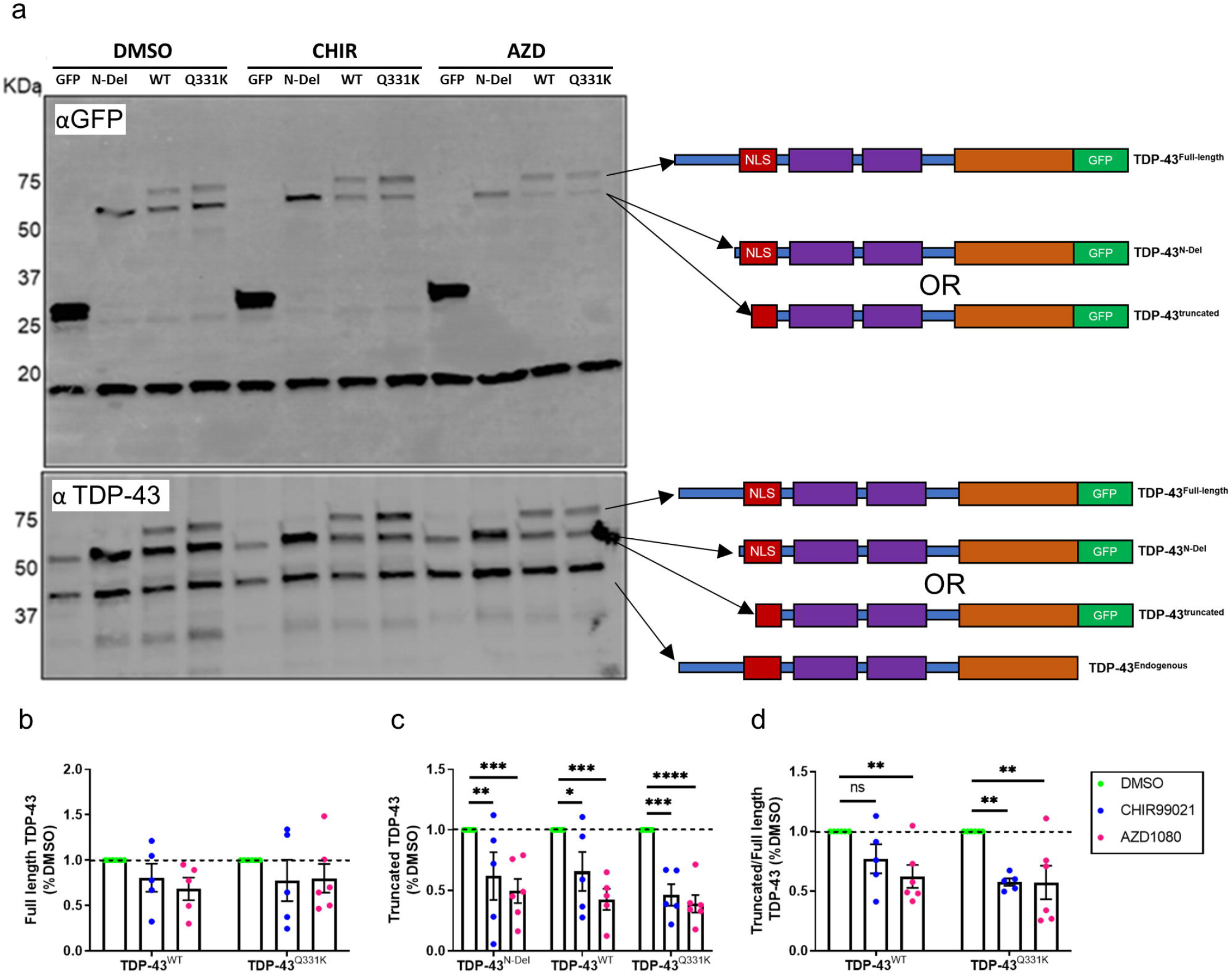
GSK-3 inhibition preferentially reduces the abundance of truncated TDP-43. **a.** Representative immunoblot of SH-SY5Y cells transfected with expression constructs for GFP and GFP tagged: TDP-43^N-Del^, TDP-43^WT^ or TDP-43^Q331K^ for 24 hours and treated with the GSK3 inhibitors CHIR99021 (CHIR) or AZD1080 (AZD). **b-d.** Immunoblot band intensity quantifications showing the abundance of full length, truncated and the ratio truncated:full length TDP-43 following treatment with CHIR or AZD. **b.** Full length TDP-43, ANOVA treatments ns *P* = 0.1216. **c.** Truncated TDP-43, pairwise comparisons: TDP-43^N-Del^, CHIR vs DMSO: \*\**P* = 0.0077; AZD vs DMSO: \*\*\**P* = 0.0007; TDP-43^WT^, CHIR vs DMSO: \**P* = 0.0156; AZD vs DMSO: \*\*\**P*= 0.0003; TDP-43^Q331K^, CHIR vs DMSO: \*\*\**P* = 0.0003; AZD vs DMSO: \*\*\*\**P* < 0.0001. **d.** Ratio truncated:full length TDP-43, pairwise comparisons: TDP-43^WT^, CHIR vs DMSO: ns *P*= 0.0752; AZD vs DMSO: \*\**P* = 0.0071; TDP-43^Q331K^, CHIR vs DMSO: \*\**P*= 0.0024; AZD vs DMSO: \*\**P*= 0.0024. For **(b-d)** (*n* = 5-6 biological replicates per condition); two-way ANOVA followed by Holm-Sidak *post-hoc* test for pairwise comparisons. Error bars denote mean ± s.e.m.

### GSK3 inhibition reduces the level of nuclear TDP-43 in a caspase-dependent manner

Cytoplasmic mislocalisation and nuclear depletion of TDP-43 are hallmarks of TDP-43 proteinopathies, at least at end-stage disease. To investigate the effects of GSK3 inhibition on TDP-43 localisation, primary rat cortical neurons were transfected with GFP-tagged TDP-43 constructs (either TDP-43^WT^ or ALS-linked mutant TDP-43^A315T^). Neurons were subsequently treated with CHIR99021 in doses ranging from 0.1μM to 10μM. TDP-43-GFP intensity in the cytoplasm and nucleus was determined by automated high-content fluorescent microscopy, using a vital nuclear dye (Hoechst) as reference for the nuclear compartment, and a diffusely localised cellular marker (mApple) to outline the neuronal cytoplasm[40]. GSK3 inhibition by CHIR99021 significantly reduced the nuclear abundance of both TDP-43^WT^ and TDP-43^A315T^ in a dosedependent manner (Fig. 3a). The effect on cytoplasmic TDP-43 was less pronounced where by higher doses of the GSK3 inhibitor significantly reduced the abundance of TDP-43^WT^ but not TDP-43^A315T^ (Fig. 3b), thereby causing a reduction in the nuclear to cytoplasmic ratio of both TDP-43^WT^ and TDP-43^A315T^ (Fig. 3c). Given that the vast majority of TDP-43 is localised to the nucleus (Fig. 3a,b), inhibition of GSK3 effectively reduces the abundance of total cellular TDP-43^WT^ and TDP-43^A315T^ in a dose-dependent manner.

**Fig.3.**
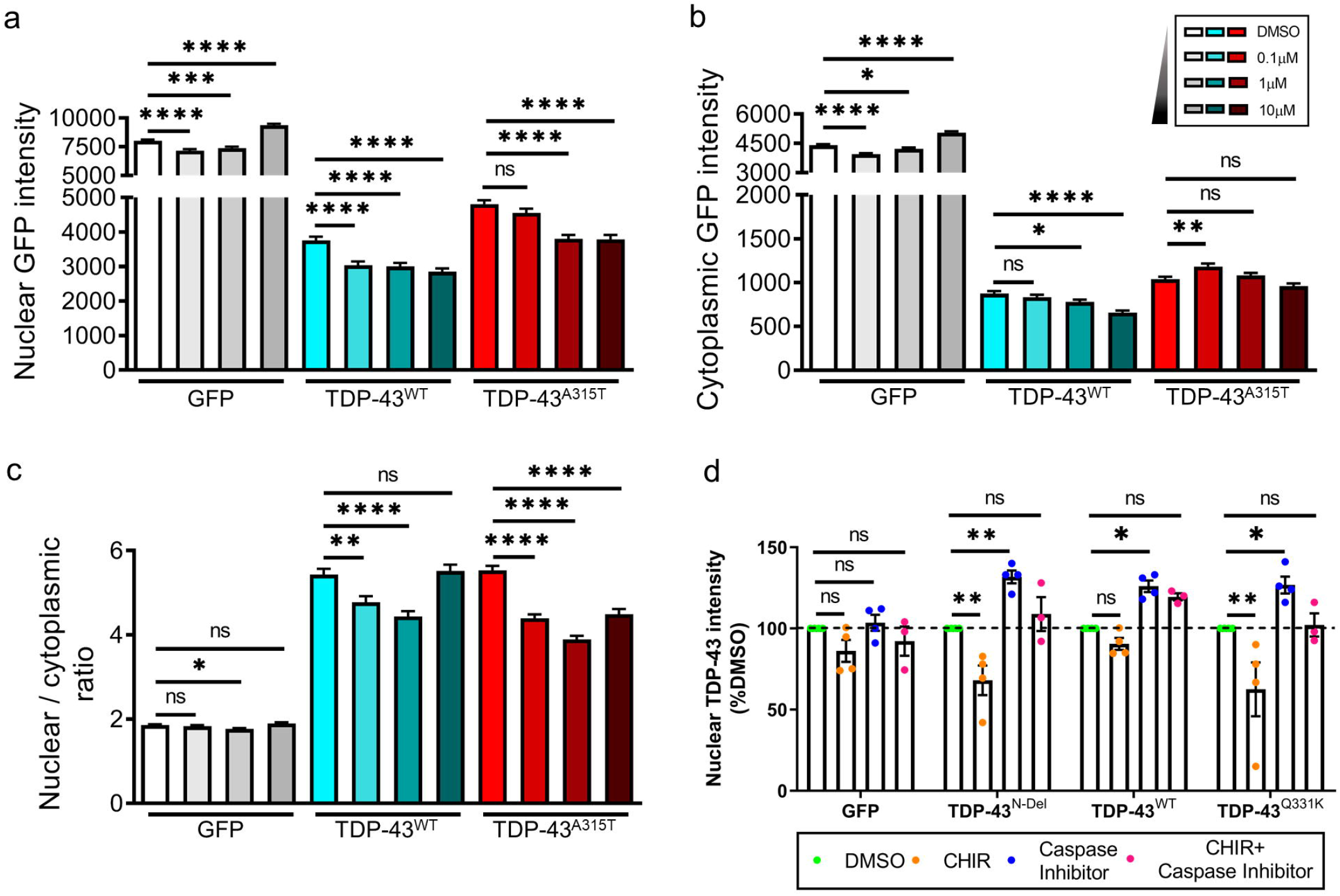
Inhibition of GSK3 reduces nuclear TDP-43 abundance in a caspase-dependant mechanism. **a-c.** Subcellular distribution and abundance of GFP and GFP tagged TDP-43^WT^ and TDP-43^A315T^ following treatment with increasing doses of the GSK3 inhibitor CHIR99021 in cortical neurons. **a.** Nuclear TDP-43, pairwise comparisons: GFP, 1μm CHIR: \*\*\**P*= 0.0003; TDP-43^A315T^, 0.1μm CHIR: ns *P*= 0.1132. **b.** Cytoplasmic TDP-43, pairwise comparisons: GFP, 1μm CHIR: \**P* = 0.0256; TDP-43^WT^, 0.1μm CHIR: ns *P*= 0.2529; 1μm CHIR: \**P*= 0.0292; TDP-43^A315T^, 0.1μm CHIR: \*\**P*=0.0023; 1μm CHIR: ns *P*=0.3181; 10μm CHIR: ns *P*=0.1379. **c.** Ratio nuclear:cytoplasmic TDP-43, pairwise comparisons: GFP, 0.1μm CHIR: ns *P*= 0.4525; 1μm CHIR: \**P* = 0.0110; 10μm CHIR: ns *P* = 0.3138; TDP-43^WT^, 0.1μm CHIR: \*\**P*=0.0018; 10μm CHIR: ns *P* = 0.6625. For **(a-c)** (*n* = 619-991 cells per condition from 3 technical replicate experiments); ****P < 0.0001; one-way ANOVA followed by Holm-Sidak *post-hoc* test for pairwise comparisons. **d.** Nuclear abundance of TDP-43 in SH-SY5Y cells transfected with expression constructs for GFP and GFP tagged: TDP-43^N-Del^, TDP-43^WT^ or TDP-43^Q331K^ for 24 hours. Cells are treated with the GSK3 inhibitor CHIR99021 and pan caspase inhibitor Q-VD-OPh. Pairwise comparisons: GFP, CHIR: ns *P*= 0.4423; Caspase inhibitor: ns *P*= 0.9809; CHIR + caspase inhibitor; ns *P* = 0.8586; TDP-43^N-Del^, CHIR: \*\**P*= 0.0062; Caspase inhibitor: \*\**P*= 0.0065; CHIR + caspase inhibitor; ns *P* = 0.8069; TDP-43^WT^, CHIR: ns *P*= 0.7312; Caspase inhibitor: \**P*= 0.0341; CHIR + caspase inhibitor; ns *P*= 0.2179; TDP-43^Q331K^, CHIR: \*\**P*= 0.0010; Caspase inhibitor: \**P*= 0.0278; CHIR + caspase inhibitor; ns *P*= 0.9962; (*n* = 3-4 biological replicates per condition); two-way ANOVA followed by Holm-Sidak *post-hoc* test for pairwise comparisons. Error bars denote mean ± s.e.m.

As both GSK3 and caspase inhibition influences the abundance of N-terminally truncated TDP-43 (Fig.1d,e and Fig. 2a,c), we hypothesised that caspase-cleavage and consequent disruption of the nuclear localisation sequence are key steps in the mechanism by which GSK3 regulates total cellular TDP-43. To test this hypothesis, we combined GSK3 inhibition with blockade of caspase activity. If N-terminal caspase cleavage occurs upstream of GSK3 mediated regulation of TDP-43, GSK3 inhibition will not alter TDP-43 expression in the absence of caspase activity. Neuroblastoma cells were transfected with constructs expressing GFP-tagged TDP-43^N-Del^, TDP-43^WT^ or TDP-43^Q331K^ and treated with either a GSK3 inhibitor, a pan caspase inhibitor or both in combination. We found that while inhibition of GSK3 reduced the nuclear abundance of TDP-43^N-Del^ and TDP-43^Q331k^ caspase inhibition had the opposite effect, causing an increase in nuclear TDP-43^N-Del^, TDP-43^WT^ and TDP-43^Q331K^ (Fig. 3.d). The ability of GSK3 inhibition to reduce the nuclear abundance of mutant TDP-43 was blocked when treated in combination with a caspase inhibitor (Fig. 3d). This indicates that GSK3 regulates the abundance of TDP-43 through an N-terminal caspase cleavage-dependent mechanism. Interestingly, the nuclear abundance of TDP-43^N-Del^ was also regulated in a similar manner to TDP-43^Q331K^. Although TDP-43^N-Del^ lacks much of the N-terminus, it retains the nuclear localisation sequence and caspase site. Thus, TDP-43^N-Del^ is still a target for caspase-mediated cleavage following GSK3 inhibition and its levels would fall in a similar fashion to that of TDP-43^WT^ and TDP-43^Q331K^ (Fig. 2a,c).

### GSK3 inhibition ameliorates TDP-43 toxicity

We previously found that loss of GSK3 suppressed TDP-43-mediated neurodegeneration in *Drosophila melanogaster[34].* To determine the therapeutic potential of targeting GSK3 in mammals, we expressed GFP tagged TDP-43^WT^ or TDP-43^A315T^ in primary rat cortical neurons and treated them with CHIR99021. The viability of single transfected cells was tracked over time using robotic microscopy[41]. Expression of both TDP-43^WT^ and TDP-43^A315T^ significantly increased the cumulative risk of death relative to expression of GFP alone, increasing the hazard ratios by 2.0 and 2.2 respectively (Fig. 4a-c). We found that CHIR99021 reduced the risk of TDP-43-mediated neuronal death in a dose-dependent manner in those cells transfected with either TDP-43^WT^ or TDP-43^A315T^ (Fig. 4a-c). Similar results were obtained in mouse primary motor neurons expressing TDP-43^N-Del^ and TDP-43^Q331K^ (Fig. 4d) using an independent luciferase-based survival assay. As GSK3 inhibition can regulate the expression of TDP-43^N-Del^ in the same manner as full length TDP-43 (Fig. 2a,c and Fig. 3d), our results suggest that TDP-43^N-Del^ is processed in the same manner as full length variants. Inhibition of GSK3 improved motor neuron survival in cells expressing N-terminally truncated TDP-43 suggesting this truncated variant exerts a modest degree of toxicity to primary motor neurons despite its inability to dimerise and fully function. TDP-43^Q331K^ was significantly more toxic to primary mouse motor neurons than TDP-43^N-Del^ and addition of CHIR99021 significantly increased the survival of motor neurons expressing TDP-43^Q331K^.

**Fig.4.**
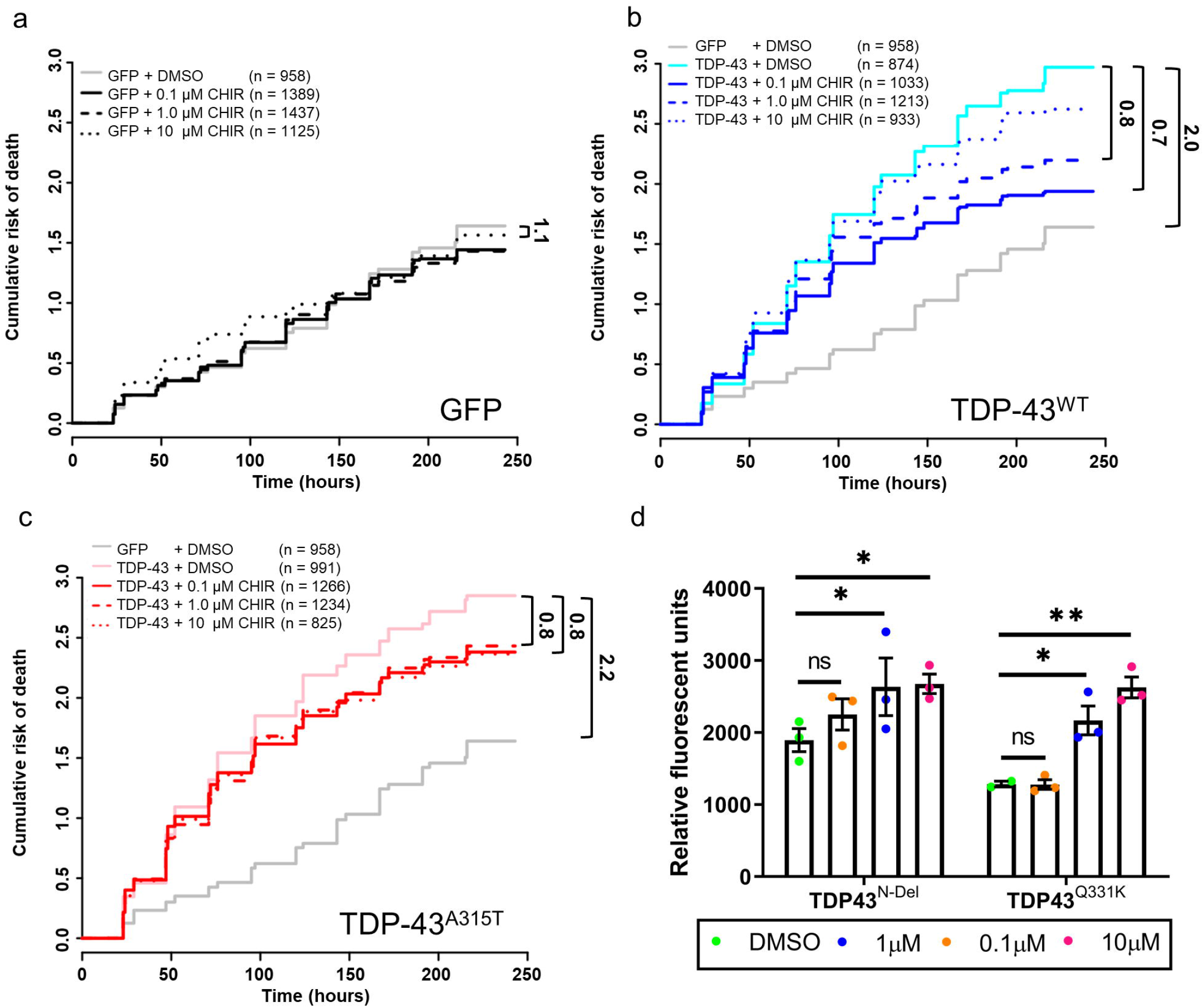
Small molecule inhibition of GSK3 ameliorates TDP-43 toxicity. **a-c.** Cumulative risk of death of primary rat cortical neurons expressing either GFP or GFP tagged TDP-43^WT^ and TDP-43^A315T^ treated with DMSO vehicle control or increasing doses of the GSK3 inhibitor CHIR99021 by longitudinal fluorescence microscopy. A hazard ratio above 1.0 indicates an increased risk of death while a value below 1.0 indicates the opposite. **a.** GFP expressing neurons, significant hazard ratios: GFP vs GFP + 0.1μM CHIR = 1.1 **b.** TDP-43^WT^ expressing neurons, significant hazard ratios: GFP vs TDP-43^WT^ = 2.0; TDP-43^WT^ vs TDP-43^WT^ + 0.1μM CHIR = 0.7; TDP-43^WT^ vs TDP-43^WT^ + 1.0μM CHIR = 0.8 **c.** TDP-43^A315T^ expressing neurons, significant hazard ratios: GFP vs TDP-43^A315T^ = 2.2; TDP-43^A315T^ vs TDP-43^A315T^ + 0.1μM CHIR = 0.8; TDP-43^A315T^ vs TDP-43^A315T^ + 1.0μM CHIR = 0.8. The number of individual neurons tracked for risk of death analysis are displayed. **d.** Survival of primary mouse motor neurons expressing either TDP-43^N-Del^ or TDP-43^Q331K^ treated with increasing doses of CHIR. Pairwise comparisons: TDP-43^N-Del^, 0.1μM CHIR: ns *P* = 0.2381; 1.0μM CHIR: \**P* = 0.0492; 10μM CHIR: \**P*= 0.0492; Pairwise comparisons: TDP-43^Q331K^, 0.1μM CHIR: ns *P*= 0.9825; 1.0μM CHIR: \**P* = 0.0312; 10μM CHIR: \*\**P* = 0.0026; (*n* = 3 biological replicates per condition); two-way ANOVA followed by Holm-Sidak *post-hoc* test for pairwise comparisons. Error bars denote mean ± s.e.m.

## Discussion

### Inhibition of GSK3 promotes neuronal survival and decreases the abundance of TDP-43 in a caspase-dependent manner

Here, we have shown that inhibition of GSK3 enhances the survival of both cortical and motor neurons expressing TDP-43^WT^ or the ALS-linked mutants TDP-43^Q331K^ and TDP-43^A315T^. In addition to its survival promoting effects, inhibition of GSK3 reduces the cellular level of caspase-cleaved TDP-43 C-terminal fragments. Caspase activity is a key requirement for both the reduction in TDP-43 abundance and survival promoting effect of GSK3 inhibitors. This suggests that GSK3 inhibitors act to enhance the turnover of TDP-43 through a caspase dependent mechanism. The result is a reduction in TDP-43 levels, which promotes neuronal survival, an observation that is in keeping with our previous finding in *Drosophila*[34]. The results we present here highlight an intriguing facet of TDP-43 cleavage, which is that the production of C-terminal fragments may have beneficial consequences. Rather than being a toxic process, cleavage of TDP-43 by caspases can cause a reduction in the abundance of full-length TDP-43, which promotes cellular survival.

### A positive feedback loop in the TDP-43-GSK3 axis could contribute to neurodegeneration

In support of our findings, a growing body of evidence indicates that inhibition of GSK3 is neuroprotective. GSK3 inhibition significantly delays disease onset and prolongs lifespan in the SOD1^G93A^ mouse model of ALS[42–44] and the GSK3 inhibitor kenpaullone prolongs survival of human pluripotent stem cell-derived motor neurons harbouring the SOD1^L144F^ or TDP-43^M337V^ mutations[45]. Chronic inhibition of GSK3 by lithium is neuroprotective against kainate-induced excitotoxic motor neuron death in organotypic slice cultures[46]. Ghrelin, a circulating hormone produced by enteroendocrine cells, protects spinal motor neurons against glutamate-induced excitotoxicity in part through PI3K/Akt-mediated inactivation of GSK3β[47]. Inhibitors of GSK3 abrogate accumulation of C-terminal TDP-43 fragments in transfected cells[33], and protect motor neurons from neuroinflammation-induced degeneration[48]. Thus, GSK3 is an attractive target for therapeutic intervention in TDP-43 linked neurodegeneration.

While inhibition of GSK3 influences TDP-43 abundance, it is also notable that TDP-43 can activate GSK3[32]. Furthermore, the abundance of GSK3β is also increased in the frontal and temporal cortices of patients with ALS and concomitant cognitive impairment[30]. Expression of TDP-43 perturbs the ER-mitochondria interface by disrupting interaction between VAPB and PTPIP51 through GSK3β activation[32]. Thus, it is clear that TDP-43 and GSK3 are fundamentally linked in a reciprocal manner, with mis-regulation of one impacting the function of the other. This intimate relationship raises the possibility that elevated TDP-43 could act in a positive feedback loop by activating GSK3 to negatively impact its own turnover. In such a situation, the abundance of TDP-43 would increase over time as its GSK3-mediated clearance is increasingly inhibited.

### TDP-43 abundance must be tightly controlled for cellular viability

TDP-43 binds a large proportion of the transcriptome and regulates several key steps of RNA metabolism[2, 49–53]. Minor alterations in TDP-43 abundance cause widespread transcriptomic changes that impact cellular function, so it is critical that mechanisms are in place to carefully regulate TDP-43 levels[54, 55]. Indeed, the level of TDP-43 is exquisitely controlled by a process of autoregulation, disruption of which is linked with ALS-FTD[54, 56–59]. Our results indicate that disruption of the caspase-dependent GSK3-TDP-43 axis is another route by which TDP-43 levels may rise with toxic consequences. At the endoplasmic reticulum, TDP-43 has been shown to be cleaved at amino acid 174 by membrane bound caspase-4 generating a 25 kDa C-terminal fragment[24, 60]. Subsequent activation of caspase-3/7 cleaves full length TDP-43 to produce a 35 kDa fragment. This sequential fragmentation reduces the abundance of TDP-43, and mitigates cytotoxicity caused by TDP-43 overexpression[24]. TDP-43 overexpression initiates caspase-4 cleavage of TDP-43 before the onset of detectable ER stress and represents a physiological mechanism to control its abundance, rather than a pathological mechanism triggered by ER stressors[25]. Thus, the fact that TDP-43 levels are tightly regulated at both RNA and protein levels emphasises the importance of TDP-43 homeostasis for cellular health.

### Misregulation of TDP-43 in neurodegenerative disease

Misregulation of TDP-43 can arise in several contexts and contribute to pathological phenotypes. The ALS associated Q331K mutation perturbs TDP-43 autoregulation thereby increasing the abundance of TDP-43[54]. Patient-derived TDP-43^M337V^ neurons have increased TDP-43 expression[61] and spinal motor neurons of patients with apparently sporadic ALS have elevated *TARDBP* mRNA[62]. The untranslated regions (UTRs) of *TARDBP* contain regulatory elements responsible for transcript stability and control, and patients with ALS demonstrate an increase in the burden of rare genetic variants in these UTRs[63]. One of these variants (c.*2076G>A in two patients with ALS-FTD) was shown to result in a doubling of *TARDBP* mRNA[64]. Under the burden of excessive *TARDBP* transcription, processing of TDP-43 at the ER could potentially be overwhelmed and contribute to a toxic increase in the abundance of full length TDP-43. The finding that TDP-43 C-terminal fragments are more prevalent in the brains of ALS-FTD patients in combination with TDP-43 aggregate pathology is a strong indicator that these mechanisms of physiological TDP-43 maintenance have been overwhelmed in disease[3].

### TDP-43 fragmentation is increased in disease

Caspase activation is a feature of several ER stressors[65] including aging[66], protein misfolding[67] and aggregation[68] and is increased in both the brains and spinal cords of patients with ALS [69, 70]. Chemical induction of apoptosis, ER stress, chronic oxidative stress and D-sorbitol induced hyper osmotic pressure can all trigger the generation of 35 kDa TDP-43 fragments in a caspase dependent manner[25]. Human lymphoblastoid lines from patients harbouring *TARDBP* mutations show that mutant TDP-43 is predisposed to fragmentation[5, 15, 71] while overexpression of mutant TDP-43^A315T^ in HEK293 cells causes persistent accumulation of protease resistant TDP-43 fragments[72]. The most common genetic cause of ALS-FTD is a hexanucleotide expansion in *C9orf72.* Repeat-associated non-AUG (RAN) translation generates poly-GA toxic repeat proteins from the repeat expansion. These poly-GA repeats induce expression of caspase-3, potentially linking *C9orf72* expansion and RAN-translation with TDP-43 proteolytic processing[73]. Levels of activated caspase-3 are also increased in spinal motor neurons of ALS patients with risk-modifying polyQ expansions of *ATXN2*[74]. While the presence of C-terminal TDP-43 fragments are a clear pathological hallmark in ALS-FTD, overexpression of 35 kDa or 25 kDa TDP-43 fragments does not necessarily cause cell death[75] or neurodegeneration *in vivo*[76]. This further supports the hypothesis that caspase mediated cleavage of TDP-43 attenuates toxicity[75] and further studies are warranted to explore how this mechanism could be targeted to alleviate TDP-43 mediated neurodegeneration.

### Conclusion

Multiple avenues of disease pathogenesis, influencing *TARDBP* transcript regulation, caspase cleavage of TDP-43 and GSK3 activity, have the potential to disrupt cellular TDP-43 homeostasis. Exposure to environmental or pathological ER stressors, missense mutations that alter TDP-43 expression or a combination of several factors, over time, could lead to a gradual failure in the homeostatic maintenance of TDP-43, causing accumulation of this protein. GSK3 inhibition reduces TDP-43 abundance in a cleavage dependent manner, alleviates TDP-43 mediated neurotoxicity, and therefore represents a target for therapy in ALS-FTD.

## Materials and Methods

### Mouse breeding and maintenance

This study was conducted on tissues from wild-type C57Bl/6J mice (*Mus musculus)* and rats (Rattus norvegicus) with breeding carried out in the UK and USA. All rodent work in the UK was conducted in accordance with the United Kingdom Animals (Scientific Procedures) Act (1986) and the United Kingdom Animals (Scientific Procedures) Act (1986) Amendment Regulations 2012. Experiments in the USA were approved by the Committee on the Use and Care of Animals (UCUCA) at the University of Michigan and performed in accordance with UCUCA guidelines. Mice were housed in cages of up to five animals under a 12 h light/dark cycle and rats were housed singly in chambers equipped with environmental enrichment.

### Plasmid constructs and small molecule compounds

The GFP-tagged TDP-43 expression constructs TDP-43^WT^ and TDP-43^Q331K^ were adapted from previously generated plasmids[6] by amplification of the TDP-43 open reading frame and ligation into the pEGFP-N1 vector. TDP-43^N-Del^ was generated by deletion of the first 81 amino acids from the N-terminus of TDP-43^WT^ using the QuickChange Site-Directed Mutagenesis Kit (Agilent).

The GSK3 inhibitor CHIR99021, CAS: 252917-06-9 was obtained from Abcam (ab120890) and a 100μM stock in DMSO stored at −20°C. AZD 1080, CAS:612487-72-6 was kindly provided by Dr Richard Mead, reconstituted in DMS0 and stored at −80°C until use. The cell-permeable, irreversible caspase inhibitor Q-VD-OPh was reconstituted in DMS0 and stored at −20°C.

### SH-SY5Y cell line culture

SH-SY5Y cells were maintained in DMEM/F-12, supplemented with GlutaMAX™ (Gibco, Thermo Fisher Scientific), 10% fetal bovine serum (FBS) (Gibco, Thermo Fisher Scientific), 1% PenicillinStreptomycin (10,000 U/ml, Thermo Fisher Scientific) and maintained at 37°C in a humidified 5% CO2 incubator.

### SH-SY5Y cell transfection and treatment

For western blot experiments, cells were passaged and plated at a density of 5 x 10^4^ cells/well in 24 well plates and allowed to recover for 24h. Cells were transiently transfected with plasmid constructs expressing GFP or GFP tagged TDP-43^N-Del^, TDP-43^WT^ or TDP-43^Q331K^ with TurboFect™ Transfection Reagent (Thermo Fisher Scientific) according to the manufacturer’s protocol. For cells treated with the small molecule GSK3 inhibitors, CHIR99021 or AZD1080, and the pan caspase inhibitor, Q-VD-OPh, compound administration was made at the time of transfection. After 24h cells were lysed in RIPA buffer containing 10μg/ml protease and phosphatase inhibitor cocktail (Merck). Lysates were cleared by centrifugation and stored at −20°C until use.

For fluorescent imaging, cells were passaged and plated at a density of 1.5 x 10^4^ cells/well in CellCarrier-96 Ultra Microplates (Perkin Elmer) previously coated with Poly-DL-ornithine hydrobromide 100mg (0.5mg/ml; Sigma). After 24h in culture, cells were transiently transfected in the same manner as for western blot experiments.

### Western blot analysis

Protein lysates from SH-SY5Y cells were quantified (bicinchoninic acid protein assay, Pierce) and then electrophoresed in 4—12% Mini-PROTEAN^®^ TGX™ Precast Gels (Bio-Rad) in 1X TGS Tris/Glycine/SDS Buffer (10X TGS Buffer, Bio-Rad). Gels were wet transferred to PVDF membranes (Millipore), blocked with a 50:50 mixture of Odyssey PBS blocking buffer (Li-Cor) and PBS with 0.1% Tween-20 for 1 h at room temperature and then probed with primary antibodies against TDP-43 (ab57105), GFP(ab290), and cyclophilin (Invitrogen PA1-027A) at 4 °C overnight, diluted 1:1000 in PBS Tween-20 (Sigma). Membranes were subsequently probed for 1 hour at room temperature with the following secondary antibodies; IRDye 680RD goat anti-Mouse and IRDye 800CW Donkey anti-rabbit (LiCor). Secondary antibodies were fluorescently tagged for Odyssey imaging. Membranes were scanned on an Odyssey^®^ CLx Imaging scanner (Li-Cor) and quantified using Image Studio Lite software.

### High content Imaging

To quantify nuclear TDP-43 intensity in SH-SY5Y cells expressing GFP tagged TDP-43 constructs, cells were imaged using an Opera Phenix High Content Imaging System (Perkin Elmer). Briefly, cells grown in CellCarrier-96 Ultra Microplates were fixed with 4% Paraformaldehyde (Thermofisher) for 15 minutes at room temperature, washed in phosphate buffered saline and treated with Hoechst 33342 (2 μg/ml; Sigma) to label nuclei. Wells were imaged in confocal mode. Digital-phase contrast images of the cell body were acquired alongside fluorescent images and data were analysed using Harmony software. Cells were partitioned into nuclear and cytoplasmic subcellular regions. The intensity of TDP-43 within each subcellular region was quantified on a cell-by-cell basis and average per well data used for downstream quantification.

### Primary rat cortical neuron cell culture and transfection

Cortices from embryonic day (E)19-20 Long-Evans rat embryos were dissected and disassociated, and primary neurons were plated at a density of 6×10^5^ cells/ml in 96-well plates, as described previously[77]. At in vitro day (DIV) 4, neurons were transfected with 100ng EGFP to mark cell bodies and 50-100ng of GFP tagged TDP-43 constructs using Lipofectamine 2000 (Invitrogen 52887), as previously described[41, 78, 79]. Following transfection, cells were placed in Neurobasal Complete Media (Neurobasal (Gibco 21103-049), 1x B27, 1x Glutamax, 100 units/mL Pen Strep (Gibco 15140-122)) and incubated at 37°C in 5% CO2. For compound treatments, neuronal media was supplemented at the time of transfection with either vehicle control or the GSK3 inhibitor CHIR99021 at concentrations ranging from 0.1μM to 10μM.

### Longitudinal fluorescence microscopy and automated image analysis

Neurons were imaged as described previously[40, 80] using a Nikon Eclipse Ti inverted microscope with PerfectFocus3a 20X objective lens and an Andor Zyla4.2 (+) sCMOS camera. A Lambda XL Xenon lamp (Sutter) with 5 mm liquid light guide (Sutter) was used to illuminate samples, and custom scripts written in Beanshell for use in μManager controlled all stage movements, shutters, and filters. Custom ImageJ/Fiji macros and Python scripts were used to identify neurons and draw both cellular and nuclear regions of interest (ROIs) based upon size, morphology, and fluorescence intensity. Fluorescence intensity of labelled proteins was used to determine protein localisation or abundance. Custom Python scripts were used to track ROIs over time, and cell death marked a set of criteria that include rounding of the soma, loss of fluorescence and degeneration of neuritic processes[41]. For measurement of nuclear and cytoplasmic protein levels, we performed automated analysis as described[40, 81]. Briefly, Hoechst vital nuclear dye was applied immediately after transfection. Nuclear ROIs were established by automated segmentation of the DAPI channel, while cellular ROIs were identified via a similar process in the RFP channel (corresponding to mApple fluorescence). The intensity of TDP-43-GFP constructs was then measured within the nuclear and cellular ROIs of each neuron, and cytoplasmic levels calculated as the difference between the cellular and nuclear ROIs.

### Primary motor neuron culture and transfection

Primary motor neurons were isolated and cultured from embryonic day 13.5 mouse embryos as previously described[82, 83]. Briefly, lumbar spinal cords were dissected, digested with trypsin and dissociated to a single cell suspension. Primary motor neurons were isolated by density gradient centrifugation using 6% Optiprep (Sigma) and cultured on glass coverslips coated with 0.5 mg/ml poly-ornithine (Sigma) and 0.5 mg/ml laminin (Thermo Fisher Scientific). Neurons were maintained in Neurobasal/B27 medium supplemented with 2% horse serum (Sigma), and 10 ng/ml each of BDNF, CNTF, and GDNF (Peprotech) with 50% media exchanges every 3 days. Primary motor neurons were transfected by magnetofection as described[82]. Motor neurons were transfected at 2 DIV using magnetic nanobeads (NeuroMag, Oz Biosciences). Culture media was exchanged 1 hour prior to transfection with Neurobasal/B27 medium without serum. Plasmid DNA was incubated with NeuroMag in minimal essential medium (MEM) for 15 minutes, and then added dropwise to the cultures. Cells were incubated on top of a magnetic plate (Oz Biosciences) for 15 minutes and after removal of the magnet, media exchanged for complete neuronal media after 1 hour.

### Motor neuron survival assay

To quantify primary motor neuron survival, neurons were co-transfected with TDP-43 expression constructs and the pGL4.50[luc2/CMV/Hygro] luciferase reporter (Promega). After 4 DIV, luciferase expression was quantified using the Bio-Glo™ Luciferase Assay System (Promega) and a PHERAstar FS plate reader. Luciferase expression was used as a proxy for the number of surviving neurons. Assay reproducibility was confirmed by manual counting of GFP-TDP-43 positive cells.

### Statistical analyses

Statistical analyses were conducted using Prism 8.4.3 (GraphPad). For comparisons between genotypes or experimental groups, multiple *t*-tests or one-way ANOVA were used when comparing two or three groups, respectively. For comparison of means split on two independent variables, two-way ANOVA was used. Multiple comparisons by were corrected using the Holm–Sidak test. For primary rat neuron survival analysis, the open-source R survival package was used to determine hazard ratios describing the relative survival between conditions through Cox proportional hazards analysis[41]. The statistical tests used and appropriate sample sizes are provided in the relevant figure legends. All statistical comparisons are based on biological replicates unless stated otherwise.

## Author Information

Francesca Massenzio present address:

Department of Pharmacy and Biotechnology FaBiT, University of Bologna, Bologna, Italy

## Acknowledgements

We thank Dr Simon Walker at the Babraham Institute Imaging Facility and Dr. George Chennell of the Wohl Cellular Imaging Centre at the Maurice Wohl Clinical Neuroscience Institute, King’s College London for imaging assistance. We also thank Dr Richard Mead for the supply of AZD 1080 GSK3 inhibitor.

## Ethics approval

All animal experiments were performed under the UK animals (Scientific Procedures) Act 1986 Amendment Regulations 2012 or were approved by the Committee on the Use and Care of Animals (UCUCA) at the University of Michigan and performed in accordance with UCUCA guidelines.

## References

1. Burrell, J.R., et al., The frontotemporal dementia-motor neuron disease continuum. Lancet, 2016. 388(10047): p. 919–31.

2. Ling, S.C., M. Polymenidou, and D.W. Cleveland, Converging mechanisms in ALS and FTD: disrupted RNA and protein homeostasis. Neuron, 2013. 79(3): p. 416–38.

3. Neumann, M., et al., Ubiquitinated TDP-43 in frontotemporal lobar degeneration and amyotrophic lateral sclerosis. Science, 2006. 314(5796): p. 130–3.

4. Arai, T., et al., TDP-43 is a component of ubiquitin-positive tau-negative inclusions in frontotemporal lobar degeneration and amyotrophic lateral sclerosis. Biochem Biophys Res Commun, 2006. 351(3): p. 602–11.

5. Kabashi, E., et al., TARDBP mutations in individuals with sporadic and familial amyotrophic lateral sclerosis. Nat Genet, 2008. 40(5): p. 572–4.

6. Sreedharan, J., et al., TDP-43 mutations in familial and sporadic amyotrophic lateral sclerosis. Science, 2008. 319(5870): p. 1668–72.

7. Benajiba, L., et al., TARDBP mutations in motoneuron disease with frontotemporal lobar degeneration. Ann Neurol, 2009. 65(4): p. 470–3.

8. Taylor, J.P., R.H. Brown, Jr., and D.W. Cleveland, Decoding ALS: from genes to mechanism. Nature, 2016. 539(7628): p. 197–206.

9. Josephs, K.A., et al., Updated TDP-43 in Alzheimer’s disease staging scheme. Acta Neuropathol, 2016. 131(4): p. 571–85.

10. McAleese, K.E., et al., TDP-43 pathology in Alzheimer’s disease, dementia with Lewy bodies and ageing. Brain Pathol, 2017. 27(4): p. 472–479.

11. Nakashima-Yasuda, H., et al., Co-morbidity of TDP-43 proteinopathy in Lewy body related diseases. Acta Neuropathol, 2007. 114(3): p. 221–9.

12. Rayaprolu, S., et al., TARDBP mutations in Parkinson’s disease. Parkinsonism Relat Disord, 2013. 19(3): p. 312–5.

13. Schwab, C., et al., Colocalization of transactivation-responsive DNA-binding protein 43 and huntingtin in inclusions of Huntington disease. J Neuropathol Exp Neurol, 2008. 67(12): p. 1159–65.

14. Nelson, P.T., et al., Limbic-predominant age-related TDP-43 encephalopathy (LATE): consensus working group report. Brain, 2019. 142(6): p. 1503–1527.

15. Rutherford, N.J., et al., Novel mutations in TARDBP (TDP-43) in patients with familial amyotrophic lateral sclerosis. PLoS Genet, 2008. 4(9): p. e1000193.

16. Cassel, J.A., et al., Characterization of a series of 4-aminoquinolines that stimulate caspase-7 mediated cleavage of TDP-43 and inhibit its function. Biochimie, 2012. 94(9): p. 1974–81.

17. De Marco, G., et al., Reduced cellular Ca(2+) availability enhances TDP-43 cleavage by apoptotic caspases. Biochim Biophys Acta, 2014. 1843(4): p. 725–34.

18. Zhang, Y.J., et al., Progranulin mediates caspase-dependent cleavage of TAR DNA binding protein-43. J Neurosci, 2007. 27(39): p. 10530–4.

19. Yamashita, T., et al., A role for calpain-dependent cleavage of TDP-43 in amyotrophic lateral sclerosis pathology. Nat Commun, 2012. 3: p. 1307.

20. Yang, Z., et al., Dual vulnerability of TDP-43 to calpain and caspase-3 proteolysis after neurotoxic conditions and traumatic brain injury. J Cereb Blood Flow Metab, 2014. 34(9): p. 1444–52.

21. Neumann, M., Molecular neuropathology of TDP-43 proteinopathies. Int J Mol Sci, 2009. 10(1): p. 232–46.

22. Arai, T., et al., Phosphorylated and cleaved TDP-43 in ALS, FTLD and other neurodegenerative disorders and in cellular models of TDP-43 proteinopathy. Neuropathology, 2010. 30(2): p. 170–81.

23. Huang, C.C., et al., Metabolism and mis-metabolism of the neuropathological signature protein TDP-43. J Cell Sci, 2014. 127(Pt 14): p. 3024–38.

24. Li, Q., et al., The cleavage pattern of TDP-43 determines its rate of clearance and cytotoxicity. Nat Commun, 2015. 6: p. 6183.

25. Berning, B.A. and A.K. Walker, The Pathobiology of TDP-43 C-Terminal Fragments in ALS and FTLD. Front Neurosci, 2019. 13: p. 335.

26. Hergesheimer, R.C., et al., The debated toxic role of aggregated TDP-43 in amyotrophic lateral sclerosis: a resolution in sight? Brain, 2019. 142(5): p. 1176–1194.

27. Valle, C. and M.T. Carri, Which TDP-43 aggregates are toxic in ALS? Oncotarget, 2016. 7(50): p. 81973–81974.

28. Beurel, E., S.F. Grieco, and R.S. Jope, Glycogen synthase kinase-3 (GSK3): regulation, actions, and diseases. Pharmacol Ther, 2015. 148: p. 114–31.

29. Hu, J.H., et al., Protein kinase and protein phosphatase expression in amyotrophic lateral sclerosis spinal cord. J Neurochem, 2003. 85(2): p. 432–42.

30. Yang, W., C. Leystra-Lantz, and M.J. Strong, Upregulation of GSK3beta expression in frontal and temporal cortex in ALS with cognitive impairment (ALSci). Brain Res, 2008. 1196: p. 131–9.

31. Ambegaokar, S.S. and G.R. Jackson, Functional genomic screen and network analysis reveal novel modifiers of tauopathy dissociated from tau phosphorylation. Hum Mol Genet, 2011. 20(24): p. 4947–77.

32. Stoica, R., et al., ER-mitochondria associations are regulated by the VAPB-PTPIP51 interaction and are disrupted by ALS/FTD-associated TDP-43. Nat Commun, 2014. 5: p. 3996.

33. Moujalled, D., et al., Kinase Inhibitor Screening Identifies Cyclin-Dependent Kinases and Glycogen Synthase Kinase 3 as Potential Modulators of TDP-43 Cytosolic Accumulation during Cell Stress. PLoS One, 2013. 8(6): p. e67433.

34. Sreedharan, J., et al., Age-Dependent TDP-43-Mediated Motor Neuron Degeneration Requires GSK3, hat-trick, and xmas-2. Curr Biol, 2015. 25(16): p. 2130–6.

35. Chang, C.K., et al., The N-terminus of TDP-43 promotes its oligomerization and enhances DNA binding affinity. Biochem Biophys Res Commun, 2012. 425(2): p. 219–24.

36. Shiina, Y., et al., TDP-43 dimerizes in human cells in culture. Cell Mol Neurobiol, 2010. 30(4): p. 641–52.

37. Zhang, Y.J., et al., Aberrant cleavage of TDP-43 enhances aggregation and cellular toxicity. Proc Natl Acad Sci U S A, 2009. 106(18): p. 7607–12.

38. Afroz, T., et al., Functional and dynamic polymerization of the ALS-linked protein TDP-43 antagonizes its pathologic aggregation. Nat Commun, 2017. 8(1): p. 45.

39. Jiang, L.L., et al., The N-terminal dimerization is required for TDP-43 splicing activity. Sci Rep, 2017. 7(1): p. 6196.

40. Malik, A.M., et al., Matrin 3-dependent neurotoxicity is modified by nucleic acid binding and nucleocytoplasmic localization. Elife, 2018. 7.

41. Weskamp, K., et al., Monitoring Neuronal Survival via Longitudinal Fluorescence Microscopy. J Vis Exp, 2019(143).

42. Feng, H.L., et al., Combined lithium and valproate treatment delays disease onset, reduces neurological deficits and prolongs survival in an amyotrophic lateral sclerosis mouse model. Neuroscience, 2008. 155(3): p. 567–72.

43. Koh, S.H., et al., Inhibition of glycogen synthase kinase-3 suppresses the onset of symptoms and disease progression of G93A-SOD1 mouse model of ALS. Exp Neurol, 2007. 205(2): p. 336–46.

44. Ahn, S.W., et al., Neuroprotective effects of JGK-263 in transgenic SOD1-G93A mice of amyotrophic lateral sclerosis. J Neurol Sci, 2014. 340(1-2): p. 112–6.

45. Yang, Y.M., et al., A small molecule screen in stem-cell-derived motor neurons identifies a kinase inhibitor as a candidate therapeutic for ALS. Cell Stem Cell, 2013. 12(6): p. 713–26.

46. Caldero, J., et al., Lithium prevents excitotoxic cell death of motoneurons in organotypic slice cultures of spinal cord. Neuroscience, 2010. 165(4): p. 1353–69.

47. Lim, E., et al., Ghrelin protects spinal cord motoneurons against chronic glutamate-induced excitotoxicity via ERK1/2 and phosphatidylinositol-3-kinase/Akt/glycogen synthase kinase-3beta pathways. Exp Neurol, 2011. 230(1): p. 114–22.

48. Reinhardt, L., et al., Dual Inhibition of GSK3beta and CDK5 Protects the Cytoskeleton of Neurons from Neuroinflammatory-Mediated Degeneration In Vitro and In Vivo. Stem Cell Reports, 2019. 12(3): p. 502–517.

49. Tollervey, J.R., et al., Analysis of alternative splicing associated with aging and neurodegeneration in the human brain. Genome Res, 2011. 21(10): p. 1572–82.

50. Coyne, A.N., B.L. Zaepfel, and D.C. Zarnescu, Failure to Deliver and Translate-New Insights into RNA Dysregulation in ALS. Front Cell Neurosci, 2017. 11: p. 243.

51. Prasad, A., et al., Molecular Mechanisms of TDP-43 Misfolding and Pathology in Amyotrophic Lateral Sclerosis. Front Mol Neurosci, 2019. 12: p. 25.

52. Freibaum, B.D., et al., Global analysis of TDP-43 interacting proteins reveals strong association with RNA splicing and translation machinery. J Proteome Res, 2010. 9(2): p. 1104–20.

53. Blokhuis, A.M., et al., Comparative interactomics analysis of different ALS-associated proteins identifies converging molecular pathways. Acta Neuropathol, 2016. 132(2): p. 175–196.

54. White, M.A., et al., TDP-43 gains function due to perturbed autoregulation in a Tardbp knock-in mouse model of ALS-FTD. Nature Neuroscience, 2018. 21(4): p. 552–563.

55. Fratta, P., et al., Mice with endogenous TDP-43 mutations exhibit gain of splicing function and characteristics of amyotrophic lateral sclerosis. EMBO J, 2018. 37(11).

56. Avendano-Vazquez, S.E., et al., Autoregulation of TDP-43 mRNA levels involves interplay between transcription, splicing, and alternative polyA site selection. Genes Dev, 2012. 26(15): p. 1679–84.

57. Ayala, Y.M., et al., TDP-43 regulates its mRNA levels through a negative feedback loop. EMBO J, 2011. 30(2): p. 277–88.

58. Bembich, S., et al., Predominance of spliceosomal complex formation over polyadenylation site selection in TDP-43 autoregulation. Nucleic Acids Res, 2014. 42(5): p. 3362–71.

59. Polymenidou, M., et al., Long pre-mRNA depletion and RNA missplicing contribute to neuronal vulnerability from loss of TDP-43. Nat Neurosci, 2011. 14(4): p. 459–68.

60. Yin, P., et al., Caspase-4 mediates cytoplasmic accumulation of TDP-43 in the primate brains. Acta Neuropathol, 2019. 137(6): p. 919–937.

61. Bilican, B., et al., Mutant induced pluripotent stem cell lines recapitulate aspects of TDP-43 proteinopathies and reveal cell-specific vulnerability. Proc Natl Acad Sci U S A, 2012. 109(15): p. 5803–8.

62. Koyama, A., et al., Increased cytoplasmic TARDBP mRNA in affected spinal motor neurons in ALS caused by abnormal autoregulation of TDP-43. Nucleic Acids Res, 2016. 44(12): p. 5820–36.

63. Morgan, S., et al., A comprehensive analysis of rare genetic variation in amyotrophic lateral sclerosis in the UK. Brain, 2017. 140(6): p. 1611–1618.

64. Gitcho, M.A., et al., TARDBP 3’-UTR variant in autopsy-confirmed frontotemporal lobar degeneration with TDP-43 proteinopathy. Acta Neuropathol, 2009. 118(5): p. 633–45.

65. Hitomi, J., et al., Involvement of caspase-4 in endoplasmic reticulum stress-induced apoptosis and Abeta-induced cell death. J Cell Biol, 2004. 165(3): p. 347–56.

66. Martinez, G., et al., Endoplasmic reticulum proteostasis impairment in aging. Aging Cell, 2017. 16(4): p. 615–623.

67. Oakes, S.A. and F.R. Papa, The role of endoplasmic reticulum stress in human pathology. Annu Rev Pathol, 2015. 10: p. 173–94.

68. Ogen-Shtern, N., T. Ben David, and G.Z. Lederkremer, Protein aggregation and ER stress. Brain Res, 2016. 1648(Pt B): p. 658–666.

69. Atkin, J.D., et al., Endoplasmic reticulum stress and induction of the unfolded protein response in human sporadic amyotrophic lateral sclerosis. Neurobiol Dis, 2008. 30(3): p. 400–7.

70. Ilieva, E.V., et al., Oxidative and endoplasmic reticulum stress interplay in sporadic amyotrophic lateral sclerosis. Brain, 2007. 130(Pt 12): p. 3111–23.

71. Corrado, L., et al., High frequency of TARDBP gene mutations in Italian patients with amyotrophic lateral sclerosis. Hum Mutat, 2009. 30(4): p. 688–94.

72. Guo, W., et al., An ALS-associated mutation affecting TDP-43 enhances protein aggregation, fibril formation and neurotoxicity. Nat Struct Mol Biol, 2011. 18(7): p. 822–30.

73. Zhang, Y.J., et al., Aggregation-prone c9FTD/ALS poly(GA) RAN-translated proteins cause neurotoxicity by inducing ER stress. Acta Neuropathol, 2014. 128(4): p. 505–24.

74. Hart, M.P. and A.D. Gitler, ALS-associated ataxin 2 polyQ expansions enhance stress-induced caspase 3 activation and increase TDP-43 pathological modifications. J Neurosci, 2012. 32(27): p. 9133–42.

75. Suzuki, H., K. Lee, and M. Matsuoka, TDP-43-induced death is associated with altered regulation of BIM and Bcl-xL and attenuated by caspase-mediated TDP-43 cleavage. J Biol Chem, 2011. 286(15): p. 13171–83.

76. Li, Y., et al., A Drosophila model for TDP-43 proteinopathy. Proc Natl Acad Sci U S A, 2010. 107(7): p. 3169–74.

77. Saudou, F., et al., Huntingtin acts in the nucleus to induce apoptosis but death does not correlate with the formation of intranuclear inclusions. Cell, 1998. 95(1): p. 55–66.

78. Barmada, S.J., et al., Autophagy induction enhances TDP43 turnover and survival in neuronal ALS models. Nat Chem Biol, 2014. 10(8): p. 677–85.

79. Barmada, S.J., et al., Cytoplasmic mislocalization of TDP-43 is toxic to neurons and enhanced by a mutation associated with familial amyotrophic lateral sclerosis. J Neurosci, 2010. 30(2): p. 639–49.

80. Barmada, S.J., et al., Amelioration of toxicity in neuronal models of amyotrophic lateral sclerosis by hUPF1. Proc Natl Acad Sci U S A, 2015. 112(25): p. 7821–6.

81. Archbold, H.C., et al., TDP43 nuclear export and neurodegeneration in models of amyotrophic lateral sclerosis and frontotemporal dementia. Sci Rep, 2018. 8(1): p. 4606.

82. Fallini, C., G.J. Bassell, and W. Rossoll, High-efficiency transfection of cultured primary motor neurons to study protein localization, trafficking, and function. Mol Neurodegener, 2010. 5: p. 17.

83. Fallini, C., et al., The survival of motor neuron (SMN) protein interacts with the mRNA-binding protein HuD and regulates localization of poly(A) mRNA in primary motor neuron axons. J Neurosci, 2011. 31(10): p. 3914–25.

